# The elephant in the room- Universal coverage for Costly treatments in an upper middle income country

**DOI:** 10.1101/214296

**Authors:** RP Kaur, GF Ho, MY Mastura, PP Goh, Mohamad Aziz Salowi, AH Muhd. Radzi, Anwar Hau, Jameela Sathar, Z Robaayah, Benedict Selladurai, Abu Bakar Suleiman, Zaki Morad, A Ghazali, TO Lim

## Abstract

**Background:** Upper middle income countries have made substantial progress towards universal health coverage. We investigated whether the coverage extended to diseases that incur catastrophic health spending, the contribution of pooled financing and the factors driving it in Malaysia.

**Methods:** We adapted the WHO definition of catastrophic health spending to define costly treatment as one that cost, at prevailing market price, more than 10% of the median annual household income in Malaysia. Coverage is defined as the proportion of patients in a year who were in need of a treatment and who received it. Data to estimate coverage and financing were extracted from the published and grey literature, as well as secondary data sources available on disease epidemiology and healthcare in Malaysia.

**Results:** We found coverage varies from universal for dialysis, cataract surgery, medicines for organ transplant and CML, to practically none for HCV, stroke, psoriasis and epilepsy surgery. Coverage of targeted therapies for solid cancers, knee replacement surgery, anti-TNF for arthritis and coagulation factors for haemophilia were poor while iron chelation for thalassemia, coronary revacularization, epoetin and anti-retrovirals were barely adequate. Coverage correlates negatively (r=-0.82) with health benefits foregone, and is entirely driven by the contribution of pooled financing (r=0.99 p<0.0001). The relative effectiveness of a treatment, its budget impact, media coverage and political influence of the disease area have little influence on financing. Only effectiveness of the leadership representing the therapy area is influential; an increase in one point on the leadership effectiveness scale is associated with 30% increase in the contribution of pooled financing.

**Conclusion:** Coverage for catastrophically costly treatments is uneven and inequitable in Malaysia, despite most of these are affordable. Decisions on coverage are driven by political-economic consideration.

## Background

Universal health coverage (UHC) means providing all people with access to needed health services and ensuring that they are protected financially from catastrophic health spending [1].

Upper middle income countries (UMIC), which comprise 53 countries and 33% of the world’s population, have made substantial progress towards UHC in the past decades [2]. These relatively wealthy countries spent USD 1935 per capita on health, 12 times higher than low income countries (LIC) [3]. However they are usually lumped together with LIC in most discussion on health but the challenges they face are likely to differ widely.

In health financing, these countries could typically tap into multiple funding pools. Malaysia for example has a government pension scheme to cover for state employees and pensioners, mandatory social insurance financed through a payroll tax for low income employees, voluntary private health insurance or employer sponsored plan for the middle classes, and finally for everybody including the poor, the unemployed or informally employed, tax funded safety net healthcare provided through a network of public hospitals and clinics mostly operated by the Ministry of Health (MOH) as well as some by Ministry of Education and Defence, alongside supplementary funding by Zakat for Muslim indigents and other charities for non-Muslim poor [4]. Such a fragmented multi-risk pools multi-tiered health system, an inherited legacy of British colonial rule and subsequent incremental changes, is surprisingly effective in covering the entire population for basic health services such as vaccinations, common prescription medicines, out-patient visits and in-patient acute care including surgeries. For example, in Malaysia, 97% of children aged 1 year were vaccinated against Diphtheria, Tetanus and Pertussis (DTP3) [5], 98% of mothers had antenatal care coverage [5], 99% of births were attended by skilled health personnel [6]. The number of consultations with doctors was 3.5 per person per year and hospital discharge was 119 per 1000 population, both figures are comparable to many rich OECD countries [7]. On access to prescription medicines, recent data from comparative surveys [8,9] across numerous countries have shown that a wide range of essential medicines for the treatment of cardiovascular disease (aspirin, β blockers, angiotensin-converting enzyme inhibitors and statins) and asthma (beclometasone, budesonide and salbutamol) were widely available and affordable in Malaysia and other UMICs. And all these were delivered at modest financial cost to payers. The safety net healthcare in Malaysia costs the taxpayers a mere USD423 per capita in 2013[10], the benefits loss ratio for social insurance was 0.9 [11], the medical loss ratio for Private Health insurance (PHI) was only 0.75 [12], clearly a highly profitable margin (and one that would be illegal in the US [13]). Thus, Malaysia was able to claim it has accomplished UHC with a relatively low spending on health at USD 1805 per head (below UMIC average), a mere 4% of its GDP [3,10, 14,15,16].

There is however an elephant in the room, coverage for catastrophically costly but life-saving or health improving treatments, which all payers pretended aren’t necessary. And since payers are currently not required, constitutionally or contractually, to cover for them, the financial risk has to be borne by the patients in spite of them belonging to one or more of the above mentioned risk pools. Many patients have to pay directly out-of-pocket (OOP) to access care, and many more had likely forgone treatment. In 2013, OOP accounted for a staggering 84% of private health spending [10], which would have predictable adverse and inequitable health and financial consequences [17] though data on these are scarce. Recent research [18,19] however has borne this out. A study on avoidable deaths showed that of the 2500 women who died of breast cancer in Malaysia in 2012, 50% of these deaths were premature on account of lack of access to screening and treatment services [18], and unsurprisingly, the poor bore the brunt of this. Another study investigated the financial consequences of cancer and found that among patients treated at public hospitals, 45% had experienced financial catastrophe within 12 months after diagnosis, and OOP spending has pushed 51% of the households of surviving patients into economic hardship [19].

The aim of this research is to determine the coverage for a wide range of catastrophically costly treatments which are indicated for the disease burden common in Malaysia.

## Methods

This study is based on data extracted from the published and grey literature, as well as secondary data sources available on disease epidemiology and healthcare in Malaysia.

### Inclusion criteria

We adapted the WHO definition of catastrophic health spending [20] to define costly treatment as one that cost, at prevailing market price, more than 10% of the median annual household income in Malaysia. Inclusion requires data be available on the disease epidemiology and healthcare. Innovative therapies are excluded because they are unlikely to be widely accessible until decades after they are first introduced (one exception included is Direct Acting Anti-virals (DAA) for Hepatitis C infection (HCV) for which low cost generic versions are available).

### Estimation of treatment coverage

According to the Institute of Medicine [21], access is the timely use of medicine or treatment to achieve the best possible outcome. We operationalize this to define coverage as the proportion of patients in a year who were in need of a treatment (NINT) and who received it (NT).

For drug based treatments, NINT is estimated using epidemiological evidence-based approach underpinned by authoritative treatment guideline for a particular disease. Disease epidemiology data are taken from the relevant registries [22-33], published studies [34-39] or global databases [40-45]. For some procedure based treatments such as dialysis, cataract and epilepsy surgery, NINT are readily measured as these procedures are definitely indicated for all patients with End-stage renal disease (ESRD), visually impaired cataract or selected patients with refractory epilepsy respectively. The incidence of these conditions estimates the NINT. However, for other procedures such as coronary revascularization and arthroplasty, it is difficult to determine the true incidence of patients with the appropriate indication for the procedures. There is a huge variation in treatment rates even among wealthy countries [7, 46] (where presumably all who needed the treatment received it). Hence we simply use the mean rate among high-income Asian countries in the OECD [7] as the benchmark.

For drug based treatments, NT is estimated based on the volume of drug utilization data from IMS [47]. Data on mean dosing per patient are calculated based on approved prescription information or local clinical practice. We validated the method by comparing its estimate of coverage versus independent estimate from epidemiological cohort study which described the treatment uptake by patients. Data from 2 such cohorts were available, anemia in ESRD patients treated with epoetin [28] and HER2+ BC treated with trastuzumab [35]. Estimates of coverage for epoetin and trastuzumab were 63% and 26% respectively based on the drug utilization method, and the corresponding rates based on epidemiological studies were 67% and 22% (year 2012) respectively. These differences do not substantively affect the results and conclusion of this study. For procedure based treatment, NT is estimated using health services data from patient registry [28-32], and cross checked against the device utilization data for consistency.

We caution that the measurement of NT to estimate coverage should not be interpreted literally. Utilization of procedures and medicines are readily measurable. They are better interpreted as proxy for the wider range of needed services of which the medicines or procedures are only one component. For example, utilization of targeted therapies for cancer is dependent on biomarker testing.

### Estimation of Spending on treatment

Spending on a treatment is estimated by the product of NT and the unit cost per treatment for short term treatment or per year cost for longer term treatment, at prevailing market price. Spending here refers only specifically to the cost of the medicines or procedures. Other direct medical costs and non-medical costs are excluded.

### Estimation of Health benefits foregone

Health benefits foregone (HBF) is estimated using criterion-based benchmarking approach. This compares the health outcome achieved in a patient population with the recommended target outcome for a treatment [48] or the outcome reported in a reference country assumed to provide universal coverage for the treatment. Unless otherwise stated, Australia is the reference country [41, 49,50]. We caution the measurements here are at best a crude proxy of the benefits foregone as many factors, besides coverage, influence the health outcome of a patient population.

### Factors influencing universal coverage for costly treatments

Coverage for catastrophically costly treatments is expected to be largely driven by the level of pooled financing, as by definition of catastrophic care, most people in a population cannot afford to pay OOP for the treatment. We therefore investigated both non-political and political factors [51-53] that could potentially influence the level of pooled financing for the treatments included in this study. These are:

1. The NINT and the unit cost per treatment. The product of NINT and unit cost is the budget impact to payers
2. Relative effectiveness among the treatments which is the sum of the score evaluated by experts on 2 dimensions; the effect of a treatment on disease progression (curative to symptomatic relief only) and on survival outcome (live-saving treatment for rapidly fatal disease to no effect). We use expert’s judgment because there is no published data on the comparative effectiveness among these treatments, and data on the observed treatment effects are simply not available for most diseases in this population.
3. Media advocacy is measured by the count of media coverage for the disease and its treatments between 2011 and 2015 in the two largest circulating English newspaper in Malaysia.
4. Political influence on health policy-making is measured by access to sympathetic and influential policy makers in the disease or therapy area. This is scored by key informants from 1 (none) to 5 (highest influence).
5. Leadership effectiveness in the disease or therapy area. This is the sum of the score as evaluated by key informants on 2 dimensions, leadership capability and integrity. This is scored from 1 (least effective) to 5 (most effective).

## Results

Coverage varies from universal (95 to 100%) for immunosuppressive drugs for organ transplant, dialysis, cataract surgery, imatinib and nilotinib for CML, to practically none (0 to 2%) for DAAs and Interferons for HCV, alteplase for ischaemic stroke, anti-TNF for psoriasis and epilepsy surgery (Table 1). Coverage of targeted therapies for solid cancers were generally poor (7% to 24%) though hematologic cancers fared much better. Coverage for knee replacement surgery (21%), anti-TNF for Rheumatoid arthritis (26%) and prophylactic coagulation factors for Hemophilia (30%) were also poor while iron chelation for thalassemia (50%), coronary revacularization (51%), epoetin for renal anemia (63%) and anti-retrovirals (ARV) for HIV (51%) were barely adequate. Spending on these treatments varies widely from a high RM 409 million for coronary revascularization, RM 282 million for ESRD (dialysis+epoetin), RM 140 million for Knee arthroplasty and RM102 million for ARVs for HIV, to practically zero (<RM 1 million) for DAAs for HCV, alteplase for ischaemic stroke and epilepsy surgery.

**Table 1:** Coverage, Spending and Financing for Costly Treatments and Health Benefits foregone in Malaysia 2012-2015

| # | Disease | Treatment | Year | Number in Need of Treatment | Number treated | Coverage | Spending | Pooled Financing | Health benefits foregone |
| --- | --- | --- | --- | --- | --- | --- | --- | --- | --- |
|  | Unit |  |  | Per year | Per year | % | Million RM | % of spending | % of benchmark |
|  | **Cancers** |  |  |  |  |  |  |  |  |
| 1. | Stage T1c to III HER2+ BC | Trastuzumab, Lapatinib | 2012 | 1508 | 321 | 21 | 21.3 | 77 | 39 |
| 2. | Stage III & IV EGFR+ NSCLC | EGFR TKI | 2012 | 938 | 221 | 24 | 17.9 | 61 | 81 |
| 3. | Stage III & IV ALK+ NSCLC | ALK TKI Crizotinib | 2014 | 147 | 10 | 7 | 1 | 42 | 81 |
| 4. | Stage IV Colorectal CA | Bevacizumab | 2012 | 1294 | 125 | 10 | 5.5 | 53 | 34 |
| 5. | Stage III & IV Ovarian CA | Bevacizumab | 2012 | 295 | 39 | 13 | 1.73 | 53 | 46 |
| 6. | Stage IV Renal cancer | Sunitinib, Pazopanib | 2012 | 278 | 45 | 16 | 3.9 | 67 | 26 |
| 7. | B-cell NHL & CLL | Rituximab | 2012 | 1137 | 854 | 75 | 24.3 | 77 | 20 |
| 8. | Chronic Myeloid Leukemia (CML) | Imatinib | 2012 | 973 | 934 | 96 | 27.8 | 93 | 0.01 |
| 9. | Resistant CML | Nilotinib | 2012 | 300 | 292 | 97 | 10.8 | 91 |  |
| 10. | Myeloma | Bortezomib | 2012 | 209 | 94 | 45 | 5.6 | 78 | 23 |
| # | Diseases | Treatments | Year | NINT | NT | Coverage | Spending | Pooled Fin | HBF |
|  | **Non-Cancers** |  |  |  |  |  |  |  |  |
| 11. | Transplant for organ failure | MMF, CsA, Tacrolimus, Antibodies | 2012 | 2832 | 2832 | 100 | 42.2 | 83 | 7 |
| 12. | HIV CD4 count <350 cells/mm<sup>3</sup> | Anti-Retrovirals | 2014 | 42408 | 21654 | 51 | 102.0 | 99 |  |
| 13. | Chronic HCV | Sofosbuvir | 2015 | 45000 | 1 | 0 | 0.02 | 0.0001 | 99.9 |
| 14. | Chronic HCV | Interferon alfa | 2012 | 44000 | 880 | 2 | 10.9 | 78 |  |
| 15. | Ischemic Stroke | Alteplase | 2012 | 46000 | 440 | 1 | 0.5 | 72 |  |
| 16. | Moderate to Highly active RA | Anti-TNF | 2012 | 1200 | 314 | 26 | 16.0 | 83 |  |
| 17. | Severe Psoriasis | Anti-TNF | 2012 | 696 | 35 | 5 | 1.78 | 77 |  |
| 18. | Anemia in prevalent ESRD | Epoetin, target Hb>10 | 2012 | 26381 | 16596 | 63 | 107.5 | 89 | 10 |
| 19. | Transfusion dependent Thalassemia | Iron chelators (DFO, DFP, DFX), target sr. Ferritin <2500 ug/L | 2012 | 5700 | 2850 | 50 | 33.0 | 99 |  |
| 20. | Hemophilia A and | Prophylactic Factor | 2013 | 1053 | 318 | 30 | 44.8 | 99 |  |
|  | B age<18 | VIII/IX replacement |  |  |  |  |  |  |  |
|  | **Procedure based treatment** |  |  |  |  |  |  |  |  |
| 21. | Incident ESRD | Dialysis | 2012 | 6635 | 6370 | 96 | 174 | 88 | 3 |
| 22. | Coronary heart disease | Revascularization (PCI and CABG) | 2015 | 37000 | 18738 | 51 | 409.0 | 75 |  |
| 23. | Bilateral Cataract with VA<6/60 | Cataract surgery with IOL implant | 2014 | 15000 | 14292 | 95 | 42.9 | 86 | 5 |
| 24. | Refractory childhood epilepsy | Epilepsy surgery | 2014 | 945 | 12 | 1 | 0.5 | 42 | 98 |
| 25. | End stage Knee OA | Knee replacement surgery | 2015 | 34000 | 7000 | 21 | 140.0 | 62 |  |

Coverage correlates negatively (Figure 1, r=-0.82) with health benefits foregone. Low coverage is associated with poorer health outcome in the patient population compares to the recommended target or reference population.

**Figure 1:**
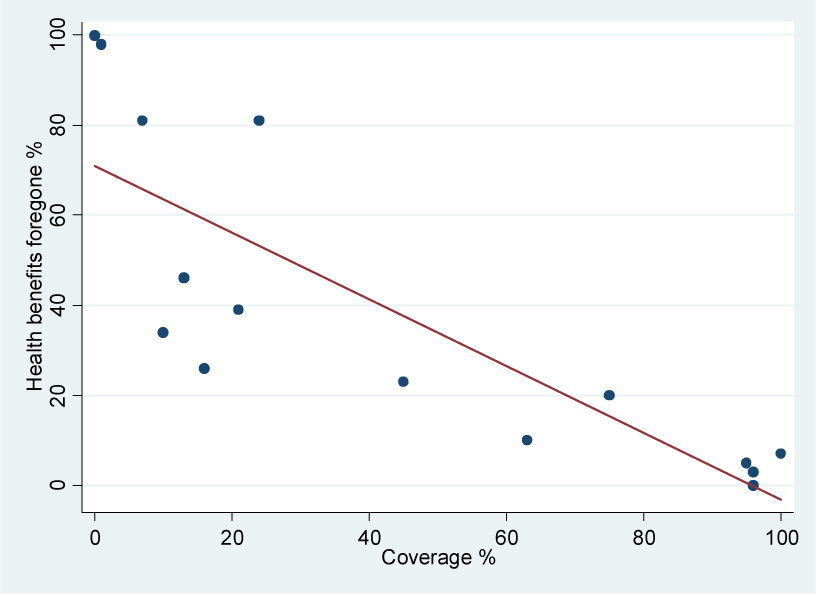
Health benefits foregone related to Treatment coverage, r= -0.82.

Coverage for costly treatment, as expected, is almost entirely driven by the contribution of pooled financing to the total required funding (Figure 2, r=0.99 p<0.0001). The contribution of pooled financing in turn may be influenced by disease, treatment and political economic factors in the country.

**Figure 2:**
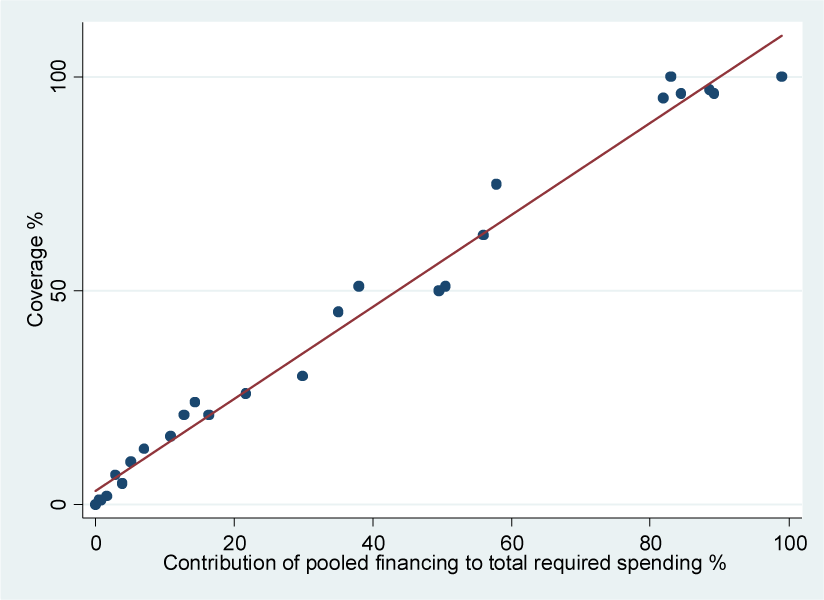
Coverage related to Contribution of pooled financing to total required funding, r=0.99.

- The relative effectiveness of a treatment, the basis of conventional health technology assessment, has practically no influence on the level of pooled financing (Figure 3, r=0.007 p=0.97).

**Figure 3:**
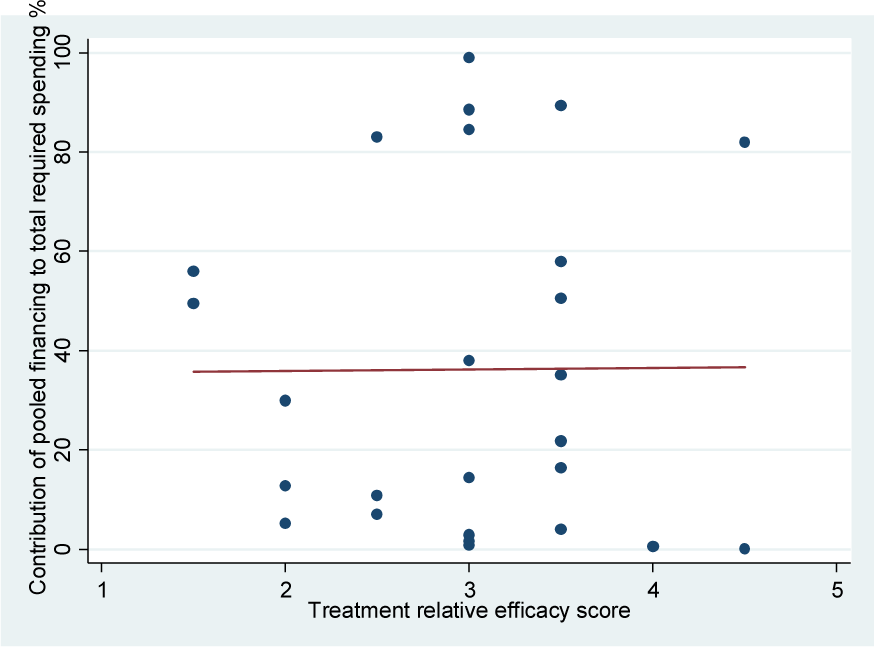
Influence of Relative Effectiveness of treatment on level of pooled financing, r= 0.007.
- The budget impact (BI) to payers as expected is negatively related to pooled financing but only modestly and statistically insignificant (Figure 4, r=-0.23, p=0.28). BI is the product of NINT and unit treatment cost, and both are also negatively correlated with the level pooled financing (r=-0.18 and r=-0.33 respectively).

**Figure 4:**
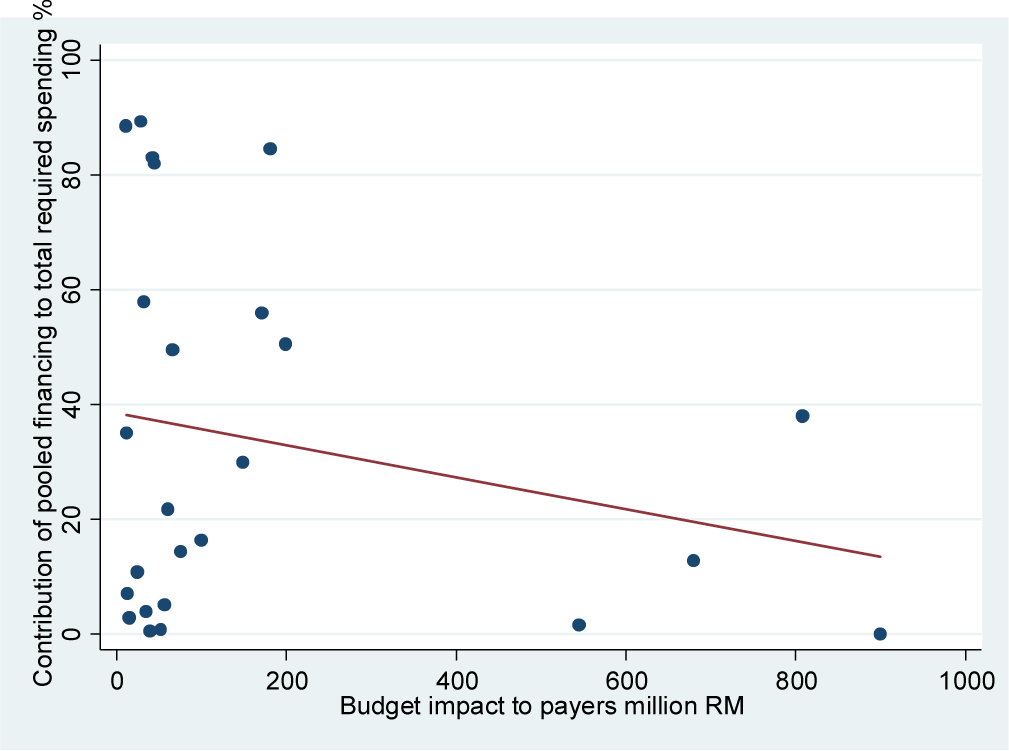
Influence of Budget impact on level of pooled financing, r= -0.23.
- Media coverage varies widely among these diseases and treatments. HIV/AIDS enjoyed the highest coverage (2366 count between 2011 and 2015), solid cancers (1402 count), kidney disease (1321), coronary heart disease (1012) and stroke (1005) were also widely covered while organ transplant, hematologic cancers, cataract and eye diseases, HCV, rheumatic and musculoskeletal diseases, psoriasis and skin diseases, epilepsy and neurological disease were poorly covered. Media coverage however has a small insignificant negative influence on pooled financing (r=-0.13 p=0.52). Likewise, the effect of political influence (r=-0.004 p=0.98).
- Leadership effectiveness has the strongest influence (Figure 5, r=0.81 p<0.0001) among all the factors investigated in this study. An increase in one point on the leadership effectiveness scale is associated with 30% increase in the contribution of pooled financing.

**Figure 5:**
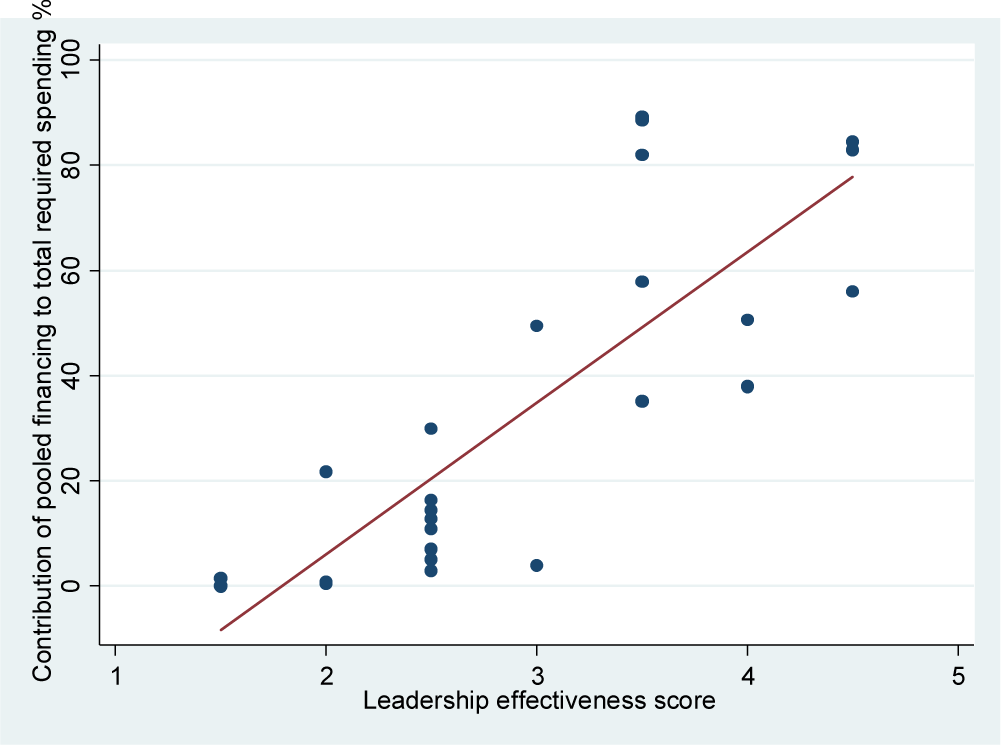
Influence of Leadership effectiveness on level of pooled financing, r=0.81.

**Figure 6:**
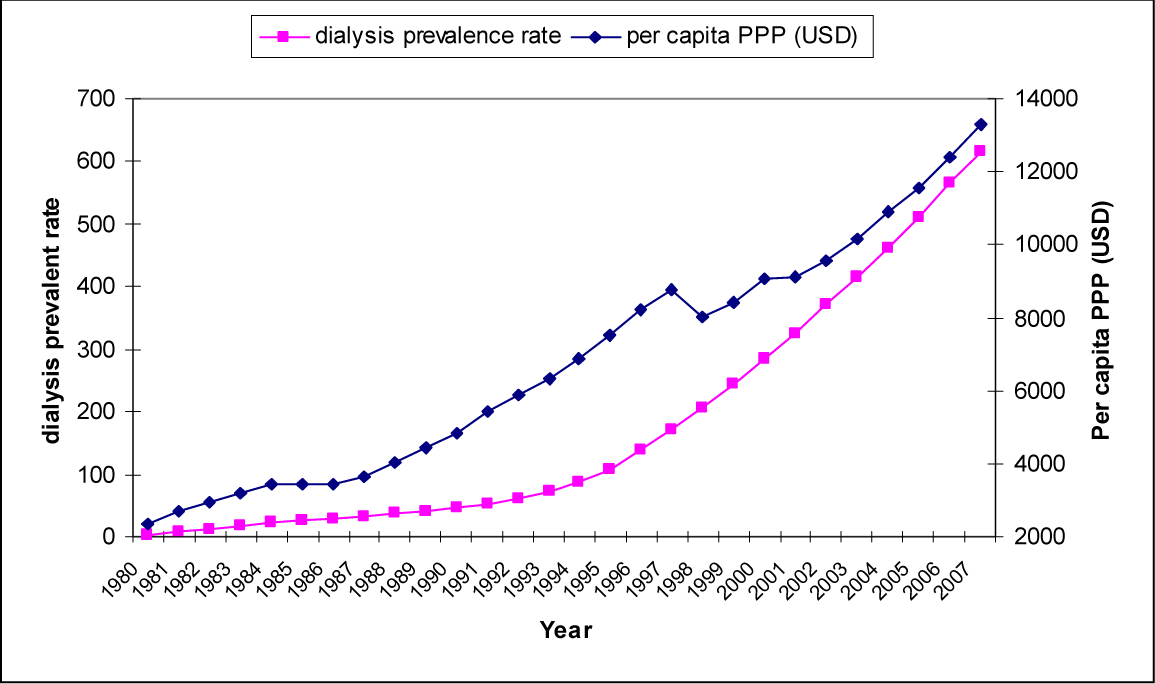
Dialysis Prevalence in Malaysia and GDP growth, 1995 to 2007.

Drug based and procedure based treatments show an interesting contrast. Drug based treatments have a higher mean unit treatment cost per patient per year compare to procedure based treatments (mean RM 44,558 versus RM22,762). Procedure based treatments however received higher level of spending (mean RM 766 million versus RM 478 million for drugs), higher level of contribution from pooled financing (53% versus 37% for drugs) and thus better mean coverage (53% versus 37% for drugs).

## Discussion

### Variation in coverage of costly treatments

We have found huge variation in spending on and coverage for costly treatments for common disease burden in Malaysia. The findings imply that a poor patient’s access to life-saving or health improving treatment is largely a function of what disease the patient has, which is a matter of luck. We have also empirically demonstrated the absolute necessity of pooled financing to ensure equitable access to costly treatments. This is consistent with the observation among countries that none could achieve universal health coverage as long as it relies predominantly on OOP for medical care [52], so likewise, within a country, no treatment could be universally covered without pooled financing.

What then explains the huge variation in coverage? There are 2 possible explanations: payers are unable to pay (affordability) or unwilling to pay.

### Payers cannot afford to pay?

Affordability is determined by cost of treatment on the one hand, and a country’s wealth and thus the resources it could mobilize for healthcare on the other. Our results show universal coverage for the treatments included here are affordable to Malaysian payers. While the total budget impact has a negative but modest effect on coverage, if the will is there, payers clearly could mobilize the necessary financial resources. They were able to provide universal coverage for immunosuppressive drugs for organ transplant, dialysis, cataract surgery, imatinib and nilotinib for CML. And some of these are not treatments with low budget impact. Dialysis and CML are two diseases with annually recurring costs (the treatments kept patients alive for relatively long duration) and hence over time they are two of the most costly treatments. The net present value of the cost of dialysis exceeded RM 1 billion a year, easily the largest budget impact by a wide margin among all treatments in the study. For these same patients, payers forked out another RM107 million in 2012 for epoetin merely to correct anemia on dialysis. To put this figure in perspective, RM107 million is equivalent to 60% of total spending on all medicines for solid cancer (cytotoxics, hormones and targeted therapies), another indication of the gross under-spending and under-coverage for solid cancer, a disease with a far higher disease burden than kidney failure.

Despite providing universal coverage for some costly treatments, the current budget impact to payers is relatively modest. The tax-funded safety net care accounted for a mere 7.25% of total government expenditure, PHI a mere 16% of private spending [10] while SOCSO’s benefit loss ratio was 0.9 (most of its benefits are for disability pension rather than healthcare) [11] with contributions from a mere 2.75% tax on payroll. These financing sources clearly have much room to grow. Improving coverage to 95% for all the treatments included in this study would cost an additional RM 920 million per year (costing assumes volume discounted prices at 75% current market prices, which is conservative based on observed inverse price-volume relationship for many therapies), a mere 2.2% of current health spending, less than what we currently spent annually on dialysis alone and well within the ability of payers to pay.

A case however may be made that the biologics and cancer medicines included in this study are relatively unaffordable to payers compare to the other treatments studied. Based on cross country comparative price studies [54,55], suppliers of these medicines have likely mispriced their products highly relative to Malaysia’s GDP at purchasing power parity. Medicine pricing in Malaysia is left entirely to the market, which has worked well for small molecule generics with many competitive suppliers [8,9]. The medicines included here are mono-sourced and the free market cannot therefore be relied upon to lower prices. In the absence of commitment to increase coverage, payers have not come forward to negotiate lower prices in exchange for larger treatment volumes. Further, to some extent, the high prices for these medicines in Malaysia are also self-inflicted. Malaysia has erected stringent regulatory barrier to generic copies of biologics. Europe for example has approved 20 biosimilars to date [56] while there are only a few such products in Malaysia. Even more non-EMA approved biologics are widely available in India, and middle income Latin American and Asian countries [56] but none of these could enter Malaysia. Malaysia’s experience with epoetin has shown that the best way to lower the price of medicines quickly and steeply is by allowing competitive supplies. Epoetin used to cost payers RM 150 per 2000IU, the introduction of alternative supplies since early 2000s has lowered its price 10 folds to a mere RM15 per 2000IU, a rare replay of Moore’s Law in health technology.

It should be noted this study has deliberately excluded innovative therapies. The ongoing debate over high-priced innovative medicines [57] is completely irrelevant outside developed countries; such medicines will never become widely accessible until many decades later in developing countries. Hence, most of the medicines included in this study are decades old and some are even obsolete, superseded by newer treatments. Epoetin for example has been available since the 1980s and trastuzumab was first approved by FDA in 1998. Both these medicines are not universally accessible yet by 2012 as shown in this study. The medical indigents are so routinely denied access to life-saving albeit costly treatments it has not struck anyone as odd that such old or even obsolete treatments should remain out of reach to the poor decades after they were first introduced.

### Payers unwilling to pay?

Given Malaysia’s demonstrable ability to pay, why coverage has yet to improve for so many treatments despite Malaysia’s UMIC status. Even in UMIC, coverage for costly treatments increases incrementally over time, if it ever did, in parallel with economic growth. This has been clearly demonstrated for dialysis as shown in Figure 10 [4]. This raises the question: why some diseases were able to capture the surplus resources made available from economic growth to fund its treatment but not others? What drives funding for a medical therapy, which in turn drives access and thus explain the variation in coverage as found in this study?

Data are scarce in developing countries. Health policy making is rarely informed by any objective data. The results reported here are the first quantitative description of the gaps in coverage for a wide range of treatments in an UMIC. We have shown in this study that the pattern of spending and coverage are not based on consideration of treatment needs or efficacy. Even economics has only a modest effect. The wide disparity in treatment coverage observed in this study can only be understood in the context of Malaysia’s political-economy. Malaysia is a relatively autocratic country where the same political party has always governed since independence 60 years ago. Public policy has long been dominated by elites in the government, bureaucracy and related interest groups. These elites are widely perceived as competent if not necessarily honest, having consistently delivered on continuing economic growth and rising living standards the past 6 decades, as well as provided for redistributive basic health, education and other social services. There is little overt public dis-affectation and the same political party has won every one of the past 14 elections with huge majorities.

In such an environment, public policy decisions are made exclusively by elites behind closed-door, though undoubtedly, depending on the issue, they are occasionally susceptible to populist pressure. The public voices on healthcare issues are not just muted but ineffectual. Catastrophic disease is not a single disease; cancers alone are hundreds of diseases. Patients, and the scores of civil society organizations and the leaderships representing them, are highly fragmented and easy to ignore. Malaysian society is also riven along ethnic-religious lines and no social movement that cut across these divides has successfully emerged on any issue since independence. Healthcare has never even made it as an electoral issue in the past 14 elections, let alone to galvanize the social forces to exert domestic pressure for change. Those same fault lines have also engendered little social solidarity in Malaysian society. The small tax-paying segment of the population appears unwilling to trust any policy that entails transferring more of their incomes to a public bureaucracy to disburse as health or other social benefits. The huge gaps in treatment coverage as described in this study highlight the underlying reality of social fragmentation in Malaysia. The upper and middle classes have long exited tax-funded safety net care for the high-quality high-priced no-waiting private care [58], and further public voices on healthcare issues [59]. The wider population, including the poorest segment who are routinely denied access to the treatments described in this study, seems to have acquiesced to such a socially fragmented tiered system, which has not generated any social protest to date.

Thus, in Malaysia, there is at present little political mileage to extending coverage to more costly treatments. Universal coverage for basic health services affects the entire population, the public seems satisfied with and appreciative of this, and policy makers are complacent of this real achievement. On the other hand, each of the diseases included in this study affects no more than a few thousands a year, and exerts even less populist pressure. This also explains why the 2 political factors investigated in this study, media advocacy and political influence, have little effect on healthcare financing

In the absence of compelling motivation for policy makers to act, we are not optimistic that the healthcare concern described in this study will be addressed, let alone be prioritized. We will just muddle along, allowing time for the benefits of continuing economic growth and rising wealth to trickle down and incrementally lift the boat of healthcare coverage for everyone. The issue will remain emergent but never gaining sufficient traction to rise up the agenda. Occasionally, perhaps due to the confluence of unusually effective leadership, an unexpected focusing event and the quirk of individual political personality, a particular treatment may break through into the policy agenda. Even then, the policy process inevitably becomes tangled in bureaucratic inertia, and will require savvy and skilful leaders in the therapy area, quietly working in the corridor of power and the bureaucracy outside the public glare, pushing their issue on the agenda and negotiating the policy details for implementation. Healthcare coverage policy will incur substantial public spending and will inevitably attract rent seekers [60]. Honest public spirited leadership is needed to fend off these opportunists. The few treatments in this study which have attained universal coverage were likely borne of such fortuitous circumstances. As much as we like to think health policy, with its direct and often acute impact on the life and health of a population, ought to operate at a higher rational plane, it is nevertheless and unavoidably fraught with the randomness, quirks and caprices of the human world.

## List of abbreviations

AIDS: Acquired Immune Deficiency syndrome
ARV: Anti-retrovirals
BC: Breast cancer
BI: Budget impact
CA: Carcinoma
CABG: Coronary Artery Bypass Graft
CLL: Chronic Lymphocytic Leukemia
CML: Chronic Myeloid Leukemia
DAA: Direct Acting Anti-virals
DTP3: Diphtheria, Tetanus and Pertussis
ESRD: End-stage renal disease
FDA: Federal Drug Administration US
GDP: Gross Domestic Product
HBF: Health benefits foregone
HCV: Hepatitis C virus infection
HIV: Human Immunodeficiency Virus
LIC: Low income countries
MOH: Ministry of Health
NHL: Non-Hodgkin Lymphoma
NINT: Number in need of a treatment
NSCLC: Non-small cell lung cancer
NT: Number treated
OECD: Organisation for Economic Co-operation and Development
OOP: Out-of-pocket
PCI: Percutaneous Coronary Intervention
RA: Rheumatoid arthritis
RM: Ringit Malaysia
TNF: Tumour necrosis factor
UHC: Universal health coverage
UMIC: Upper middle income countries
USD: US Dollars
WHO: World Health organization

## Declarations

### Ethics approval and Consent

This study was based on data extracted from the published and grey literature, as well as secondary data sources available on disease epidemiology and healthcare in Malaysia. No human subjects were enrolled for this study and no personal identifiable data were used in this research. Hence no ethics clearance was required.

### Consent for Publication

Not applicable.

No human subjects were enrolled for this study and no personal identifiable data were used in this research.

### Availability of data and material

The datasets used and/or analysed during the current study are available from the corresponding author on reasonable request

## Author’s contributions

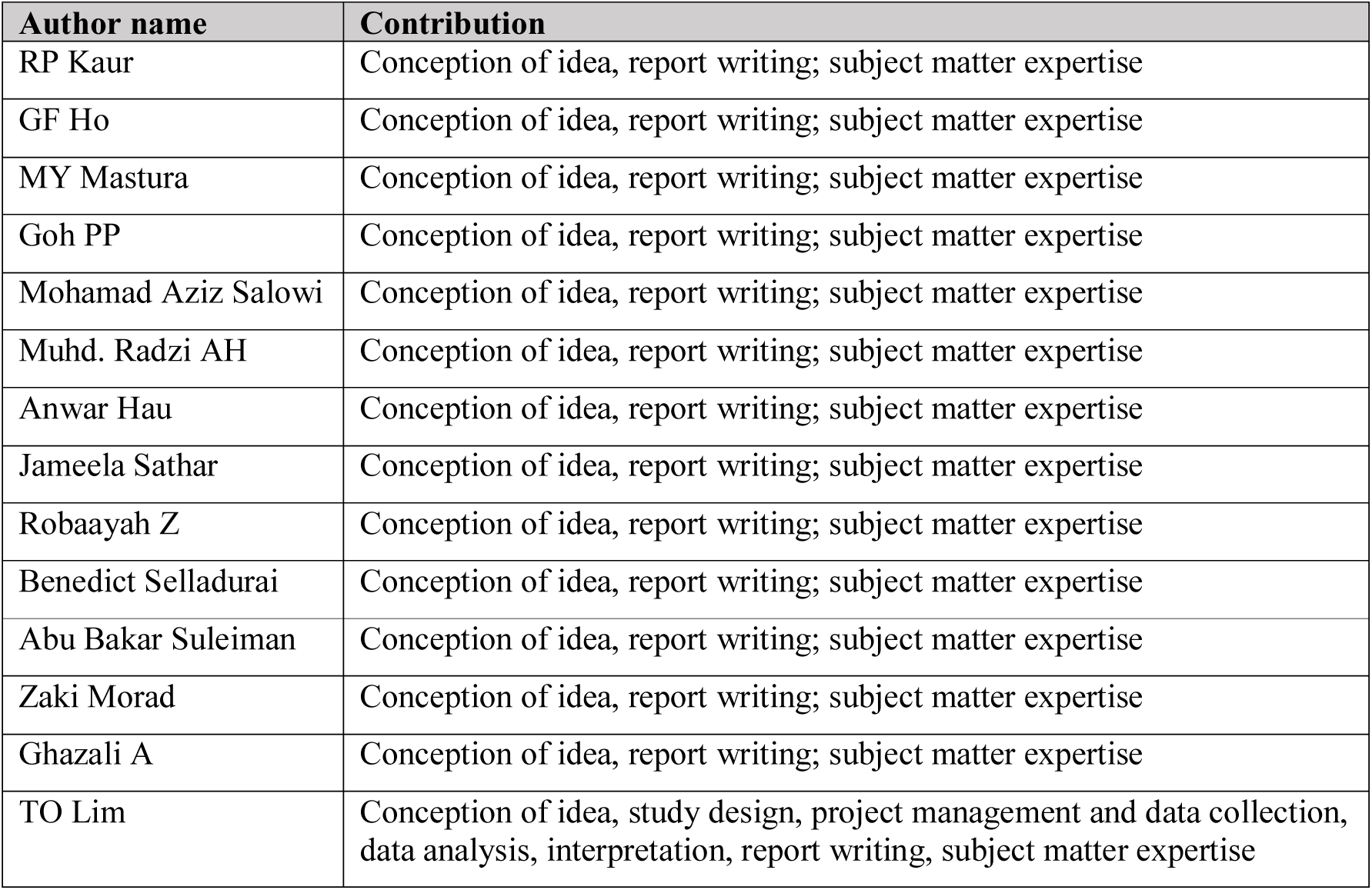

| **Author name** | **Contribution** |
| --- | --- |
| RP Kaur | Conception of idea, report writing; subject matter expertise |
| GF Ho | Conception of idea, report writing; subject matter expertise |
| MY Mastura | Conception of idea, report writing; subject matter expertise |
| Goh PP | Conception of idea, report writing; subject matter expertise |
| Mohamad Aziz Salowi | Conception of idea, report writing; subject matter expertise |
| Muhd. Radzi AH | Conception of idea, report writing; subject matter expertise |
| Anwar Hau | Conception of idea, report writing; subject matter expertise |
| Jameela Sathar | Conception of idea, report writing; subject matter expertise |
| Robaayah Z | Conception of idea, report writing; subject matter expertise |
| Benedict Selladurai | Conception of idea, report writing; subject matter expertise |
| Abu Bakar Suleiman | Conception of idea, report writing; subject matter expertise |
| Zaki Morad | Conception of idea, report writing; subject matter expertise |
| Ghazali A | Conception of idea, report writing; subject matter expertise |
| TO Lim | Conception of idea, study design, project management and data collection, data analysis, interpretation, report writing, subject matter expertise |

## Acknowledgement

The authors would like to extend their sincere gratitude and appreciation to Ms Teo JS of ClinData Consult and Ms Lena Yeap of StatsConsult for their efforts in collecting, retrieving, managing and analysing the study data. We also wish to thank all those whose names are not mentioned here who render their excellent service especially during the data collection and retrieval.

